# PKM2 ablation enhanced retinal function and survival in a preclinical model of retinitis pigmentosa

**DOI:** 10.1101/826651

**Authors:** Ethan Zhang, Joseph Ryu, Sarah R. Levi, Jin Kyun Oh, Chun Wei Hsu, Jose Ronaldo Lima de Carvalho, Xuan Cui, Tingting Yang, Stephen H. Tsang

## Abstract

Retinitis pigmentosa (RP) is a neurodegenerative disorder that causes irreversible vision loss in over 1.5 million individuals world-wide. In this study, we demonstrate that a metabolic reprogramming can treat degeneration in a *Pde6β* preclinical model of RP. Pyruvate kinase M2 (PKM2) is a glycolytic enzyme that transfers phosphate from phosphoenolpyruvate (PEP) to adenosine diphosphate (ADP), promoting glucose catabolism. Ablation of PKM2 resulted in enhanced photoreceptor survival and function in *Pde6β*-mutated mice compared with those without ablation. Electroretinogram (ERG) analyses revealed that the maximum b-wave is on average greater in *Pkm2* knockout mice than in mice with *Pkm2* intact. These rescue phenotypes from *Pkm2* ablation in a preclinical model of RP indicate that a metabolome reprogramming may be useful in treating RP.

## INTRODUCTION

Retinal degenerative diseases affect nine million Americans.^1,2^ Among these conditions, retinitis pigmentosa (RP) is the most devastating in terms of severity and prevalence, making it one of the most common cause of irreversible vision loss. Associated with more than 80 genes, RP is a heterogeneous condition that is characterized by rod photoreceptor degeneration followed by secondary cone death,^3–5^ leading to night blindness, tunnel vision, and eventually, complete blindness. Current treatment options include the use of retinal prostheses^6,7^ or *RPE65* gene therapy.^8–10^ However, these strategies are only applicable for patients with end-stage RP or those carrying *RPE65* mutations,^11^ respectively. At the same time, clinical gene therapy trials for RP primarily involve augmentation or repair of a single gene, which even if successful, would only be applicable to patients carrying those specific gene mutations.^12^ More recently, insight into the genetic etiologies of retinal degeneration has found that photoreceptor death may be linked to apoptotic factors,^13,14^ metabolic derregualation,^3,15^ and oxidative stress.^16,17^ Targeting any one of these pathways, in turn, may offer an alternative gene non-specific therapy. As of yet, however, there is no effective therapeutic option for RP and other retinal degenerative diseases.

Punzo *et al*. studied gene expression changes in RP mouse models by linking mutations in rod-specific genes to their corresponding cellular processes.^3^ Microarray analysis of samples collected at the onset of cone death revealed that 35% of annotated genes were involved in cellular metabolism, implicating metabolic processes in disease pathogenesis. To follow-up, Punzo *et al*. and Zhang *et al*. focused on the insulin/mTOR pathway and found that enhanced anabolic flux and lactate levels indeed slowed rod-cone degeneration.^18,19^ Consistently, rod-specific knockout of sirtuin 6, a transcriptional repressor of glycolytic enzymes, enhanced glycolysis, rod preservation, and photoreceptor function.^20^ Augmenting anabolism in rod photoreceptors could therefore serve as a potential treatment for RP.

One identified key regulator of cellular metabolism is pyruvate kinase M2 (PKM2), a critical catalyst for the final step of glycolysis that aids in the transfer of a phosphate group from phosphoenolpyruvate (PEP) to adenosine diphosphate (ADP). PKM2 is allosterically regulated so that it exists in two forms: an inactive dimer with low binding affinity to PEP and an active tetramer form with high affinity to PEP.^21^ When phosphorylated PKM2 is in its inactive form, less consumption of PEP occurs, resulting in higher PEP levels. This has been found to activate the pentose phosphate pathway (PPP), a process that produces glycolytic intermediates for nucleotide and lipid synthesis. As a result, when PKM2 is inhibited, subsequently, PEP accumulates, promoting the PPP.^22^ PKM2’s ability to regulate the production of glycolytic intermediates, which is essential for photoreceptor cells, makes it an attractive target to elucidate the role of glycolysis in rod and cone health. Previous studies have already found that PKM2 ablation enhances long-term photoreceptor survival in retinal detachment models, suggesting that PKM2 ablation may offer photoreceptor protection against acute external retinal stress.^23^

More recently, PKM2 has been implicated in the regulation of PDE6β,^24^ a phototransduction enzyme complex that blocks Ca+2 influx by closing cyclic nucleotide-gated (CNG) channels through cGMP hydrolysis. PDE6 is a heterotetramer with two catalytic subunits (PDE6A and PDE6B) and mutations in either can disrupt function similarly and yield identical downstream effects.^25–37^ One in 10 Americans carries a recessive RP allele,^27,38^ and 10% of those alleles confer mutations in one of the two subunits of phosphodiesterase-6 (*Pde6*), accounting for 5-8% of all RP cases.^27,28^ As such, PKM2 may very well also play a role in preserving photoreceptors from inherited retinal diseases, which represent internal sources of retinal stress.

In this report, we expand on previous findings by investigating the functional and morphological contributions of rod-specific *Pkm2* ablation in inherited retinal degeneration with a tamoxifen-inducible *Pkm2* knockout *Pde6β_H620Q/H620QRP_* model. We hypothesize that *Pkm2* knockout in the rods will promote photoreceptor anabolism and, in doing so, preserve rod and cone survival and function, delaying retinal degeneration.

## RESULTS

### Deletion of *Pkm2* in *Pde6β_H620Q/H620Q_* mice

To evaluate whether and, if so, how PKM2 plays a role in rod metabolome reprogramming, we used the tamoxifen inducible Cre recombinase system to ablate *Pkm2* in a slow progressing *Pde6β* RP model: *Pde6β*_*H620Q/H620Q*_.29 Rod selective ablation of *Pkm2* significantly reduced total PKM2 levels in *Pde6β_H620Q/H620Q_ Pkm2*_-/-_ mice compared to wild-type mice (Fig. 1).

**Figure 1.**
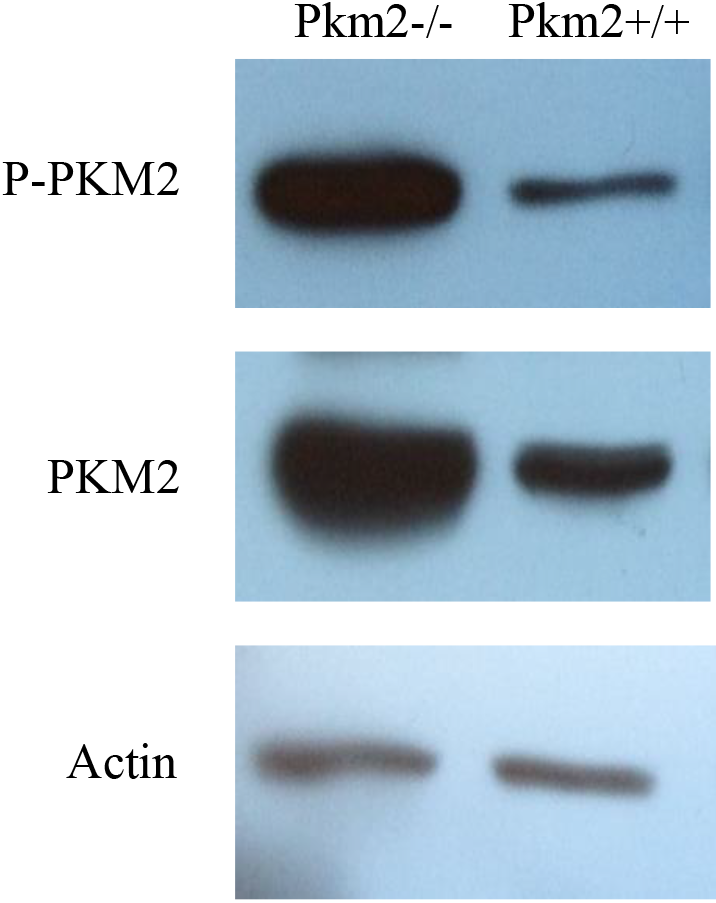
Deletion of *Pkm2* in *Pde6β_H620Q/H620Q_* mice. Immunoblot analysis was performed two weeks after tamoxifen and oil injections. Selective *Pkm2* deletion in photoreceptors reduced PKM2 and P-PKM2 expression (n=3 for both groups). Actin served as loading control.

Accordingly, there was also a reduction in phosphorylated PKM2 in *Pde6β_H620Q/H620Q_ Pkm2*_-/-_ mice, suggesting a decrease in the catalytic activity of PKM2. It is important to note, however, that experimental mice retained detectable levels of PKM2 and P-PKM2, shown in Figure 1, due to the conditional nature of the rod-specific *Pkm2* knockout.

### Deletion of *Pkm2* in *Pde6β_H620Q/H620Q_* mice improved retinal morphology

Subsequently, we gauged the survival impact of *Pkm2* ablation on retinal morphology using histology to compare the morphology of the outer nuclear layer (ONL), inner segment (IS), and outer segment (OS). *Pde6β_H620Q/H620Q_ Pkm2*_+/+_ mice exhibited degenerated photoreceptors within ONL while thin ONLs while *Pde6β_H620Q/H620Q_ Pkm2*_-/-_ mice had comparatively more cells (Fig. 2A). Similar trends were observed for IS/OS layers—compared to their isogenic controls, treated mice exhibited greater IS/OS thickness throughout the retina (Fig. 2A, B). Strikingly, all the layers in the experimental mice were still noticeably thinner than those in wild-type mice. Overall, ablation of *Pkm2* may be able to preserve but not completely rescue photoreceptor survival and preserved photoreceptors.

**Figure 2.**
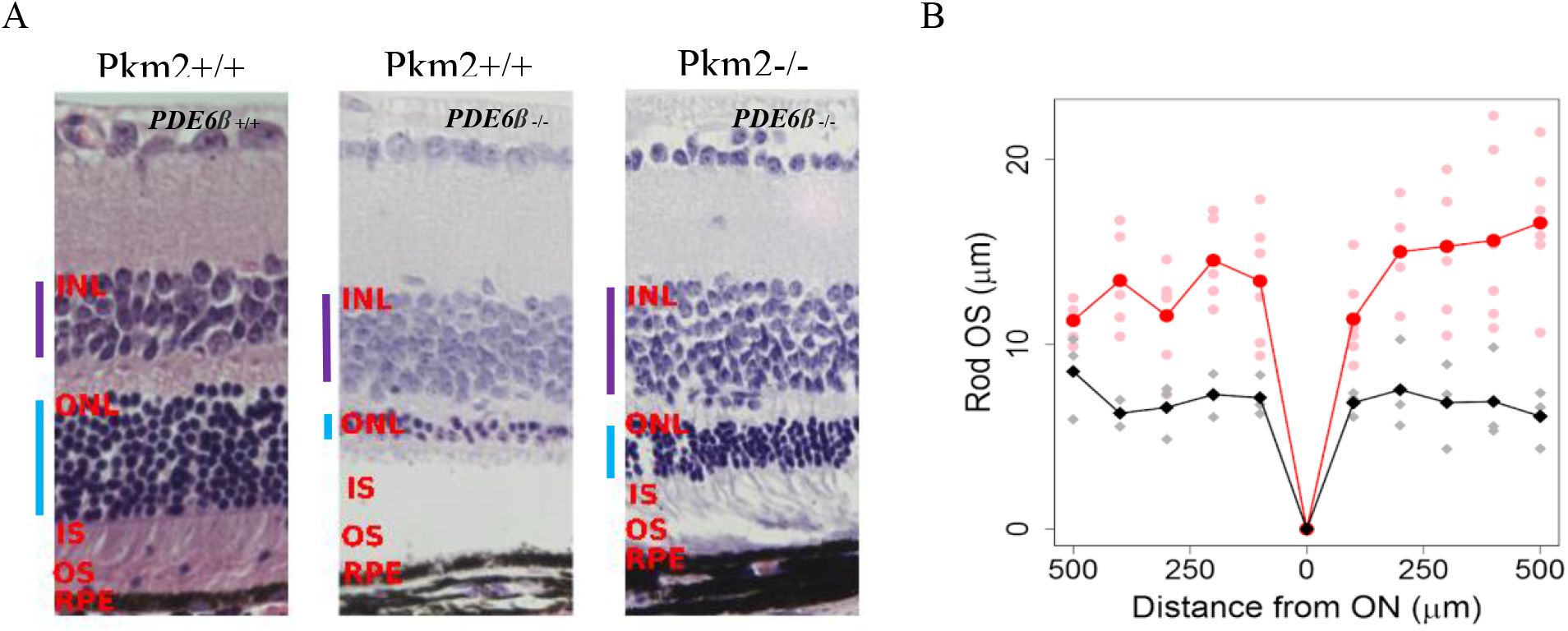
Deletion of *Pkm2* in *Pde6β_H620Q/H620Q_* mice improved retinal morphology. **(A)** H&E-stained retinal sections from were analyzed by light microscopy and compared at P60. Selective deletion of *Pkm2* enhanced but did not completely rescue ONL thickness in *Pde6β_H620Q/H620Q_ Pkm2*_-/-_ mice (n=3), but not in *Pkm2*_+/+_ *Pde6β_H620Q/H620Q_* controls. At P9, experimental mice were injected tamoxifen while wildtype and control mice were injected with oil. (Purple bar indicates INL, blue bar indicates ONL) **(B)** The difference in Rod OS length was quantified. Selection deletion of PkM2 increases Rod OS thickness at every distance from the ON (n=3). Red represents experimental mice, and black represents control mice. Highlighted line shows average Rod OS while faded dots represent individual measurements for mice.

### Deletion of *Pkm2* in *Pde6β_H620Q/H620Q_* mice improved retinal function

At four weeks post-treatment, *Pde6β_H620Q/H620Q_ Pkm2*_-/-_ mice exhibited an enhanced scotopic b-wave amplitude compared to *Pde6β_H620Q/H620Q_ Pkm2*_+/+_ (Fig. 3A), suggesting greater rod preservation. In contrast, there was no significant difference in the mixed and photopic b-wave amplitudes between the two groups, indicating that cone preservation remained approximately the same. At eight weeks post injection, all b-wave amplitudes in both groups declined considerably. Notably, *Pde6β_H620Q/H620Q_ Pkm2*_-/-_ mice maintained significantly higher scotopic, maximal, and photopic b-waves (p<0.05) compared to *Pde6β_H620Q/H620Q_ Pkm2*_-/-_ mice, indicating a more robust preservation in cone and rod function. Consistently, statistical analysis revealed a significant correlation (p<0.05) between the value of the maximal b-wave and presence of tamoxifen treatment. These results reveal that rod-specific *Pkm2* ablation improved photoreceptor vitality in *Pde6β*-mutated mice.

**Figure 3.**
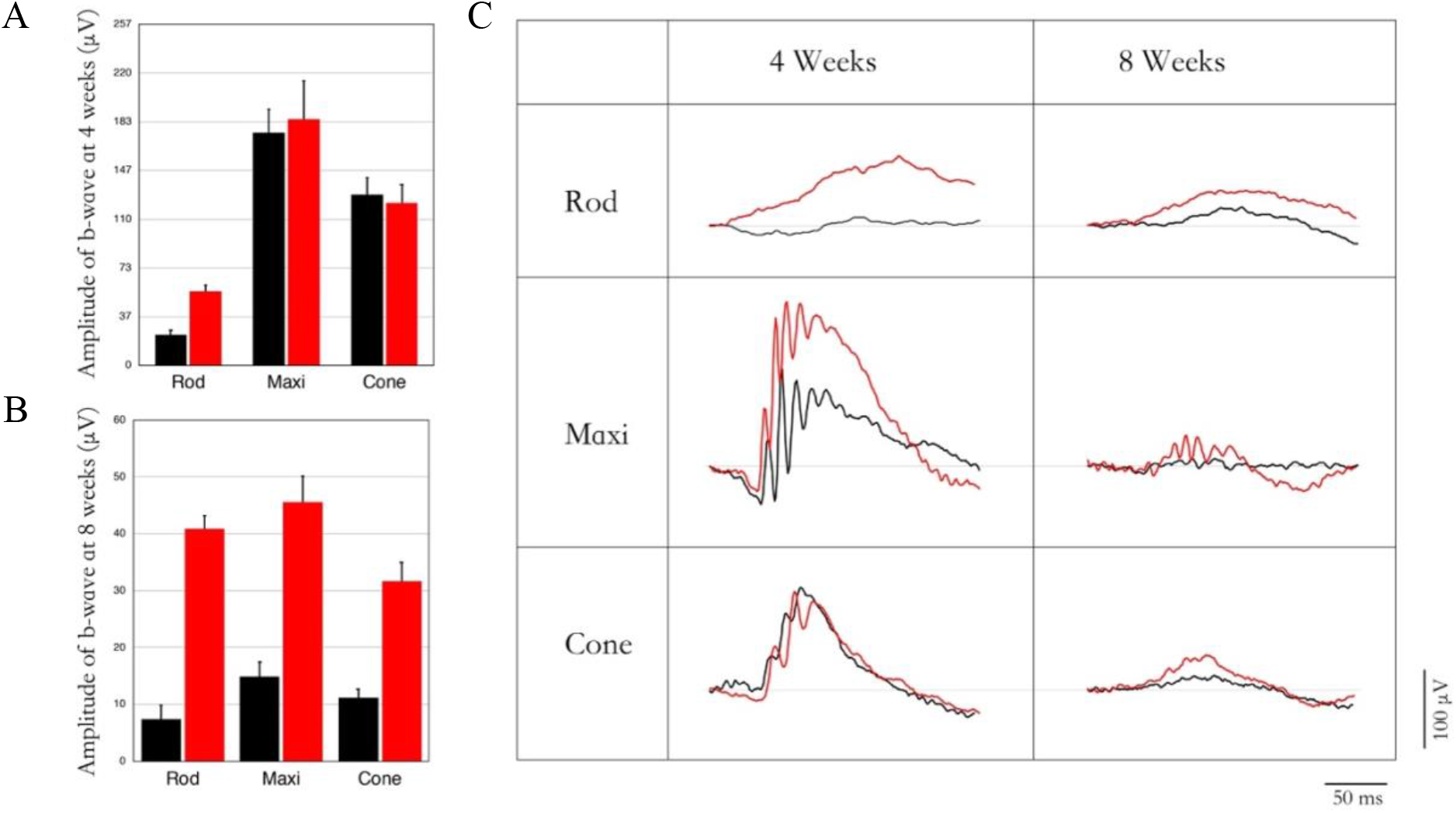
Deletion of *Pkm2* in *Pde6β_H620Q/H620Q_* mice preserves rod and cone function. ERG b-wave magnitudes for scotopic, mixed maximal, and photopic response were taken and compared in control (*Pkm2*_+/+_ *Pde6β_H620Q/H620Q_*, blue bar) and experimental *(*Pkm2*_-/-_ Pde6βH620Q/H620Q*, green bar) mice. **(A)** ERG recordings were obtained at 4 weeks post injection. Experimental mice exhibited enhanced scotopic b-wave amplitudes compared to controls. Differences in the magnitudes of mixed maximal and photopic b-waves were not statistically significant. (n=12 for both groups; (\**P* < 0.05) **(B)** ERG recordings were obtained at 8 weeks post injection. Experimental mice exhibited enhanced b-wave magnitudes for all groups compared to controls. (n=6 for control group, n=8 for experimental group; \**P* < 0.05) **(C)** Representative ERG tracings for scotopic, mixed maximal, and photopic response at 4 weeks and 8 weeks after injection.

## DISCUSSION

In this report, we demonstrated that rod-specific ablation of *Pkm2* can rescue both cone and rod photoreceptor degeneration. Using both structural and functional assays, we are the first to use a preclinical RP model to study the effects of *Pkm2* knockout. These findings further advance the work done by Chinchore *et al*. who first discovered that *Pkm2* mediates Warburg glycolysis in healthy photoreceptors, ^39^ implicating its role in photoreceptor wellbeing. Since then, various studies have sought to elucidate a direct cause-and-effect relationship. Notably, Wubben *et al*. observed that photoreceptor-specific ablation of *Pkm2* impaired photoreceptor survival and function under baseline conditions, but improved long-term photoreceptor survival under acute outer retinal stress. To account for the rescue in the retinal detachment model, Wubben *et al*. noted the upregulation of *Pkm1* and other genes involved in aerobic metabolism, proposing that accumulation of PK substrate 2-phophoenolpyruvate was sufficient to counteract the insult from iatrogenic cell damage.

Here, we advance the current understanding of PKM2 by uncovering its genetic contributions in inherited retinal degeneration. Notably, rod-specific *Pkm2* knockout noticeably increases the thickness of the ONL and IS/OS layers, indicating better photoreceptor survival. ERG functional analysis was consistent with anatomical findings and revealed that *Pkm2* ablation also improved both cone and rod function significantly. These results suggest that PkM2 deficiency was indeed able to rescue retinal degeneration. As proposed by Wubben *et al*., the rescue phenotypes may be a direct result of enhancing catabolism and therefore ATP output in photoreceptors. This explanation, however, handles photoreceptors as isolated systems whose metabolism works in a cell autonomous manner. In this paper, we propose an alternative yet complementary framework that aims to consider PKM’s role in photoreceptors in relation to the RPE, offering insight into how it may regulate the delicate metabolic dynamic between the two.

One of the first evidences of a metabolic coupling was described by Punzo, *et al*., who found that the disruptions in the insulin/mTOR signaling pathway coincided with cone death in four mouse models for RP.^3^ Strikingly, systemic supplementation of insulin was able to rescue cone death, suggesting that photoreceptor death in RP progresses in a cell non-autonomous manner. Subsequent studies characterizing the nature of this relationship have led to the emergence of the “metabolic coupling hypothesis”, which proposes that metabolic imbalances leads to cone photoreceptor death via glucose deprivation.^3,19,40^ Normally, photoreceptors take up glucose from the retinal pigment epithelium (RPE) and convert it into fat and lactate,^41^ which are subsequently used to drive essential processes throughout the retina. In photoreceptors, the steady production of protein and lipids drives the daily disposal and regeneration of the OS.^42,43^ In contrast, the RPE consumes the generated lactate as the carbon source for the tricarboxylic acid (TCA) cycle,^39,44,45^ saving glucose for photoreceptors. In RP, however, impaired photoexcitation prevents diseased rods from generating sufficient amounts of lipid and lactate. This not only stagnates OS biogenesis but also prompts the RPE to use glucose through anabolism. The depletion of glucose by the RPE, in turn, deprives photoreceptors of nourishment and leads to prevalent rod and subsequent cone death – the hallmark of late-stage RP.

In light of this framework, the rescued phenotypes indicate a rejuvenation of the disrupted metabolic dynamic within the diseased retina. Previous studies have shown that phosphorylated *Pkm2* leads to the accumulation of 2-phosphoenolpyruvate and the subsequent activation of the anabolic PPP pathway.^24^ As such, PkM2 deficiency may elevate anabolism and the synthesis of proteins and lipids, fueling the regeneration of the OS and promoting oxidative phosphorylation in the RPE. This metabolic reprogramming, in turn, restores a steady supply of glucose to the diseased photoreceptors, precluding the need to optimize metabolism. As a result, the photoreceptors return to a preferred anabolic state and employ aerobic glycolysis to satisfy their ATP needs, promoting photoreceptor health and function. It is important to note that this model is not necessarily incompatible with the notion of catabolism-mediated photoreceptor survival as described by Wubben *et al*. Roughly 80%-90% of the glucose in photoreceptors is metabolized through aerobic glycolysis, indicating that a small, but significant, fraction still undergoes catabolism. While we highlight that different anabolic processes can provide photoreceptors with both the raw materials and energy to survive and function, it is possible that a certain level of catabolism is needed to sufficiently meet the high energy demands of a healthy photoreceptor and counteract the insult from metabolic imbalance.

Another notable finding is that despite the significant improvements in both retinal morphology and function, PkM2 ablation was not able to induce a complete rescue in either case. *Pde6β_H620Q/H62Q_ Pkm2*_-/-_ mice exhibited thinner ONLs than that of wildtype mice, indicating increased photoreceptor death. Moreover, the retinal function of *Pde6β_H620Q/H62Q_ Pkm2*_-/-_ mice decreased after the initial improvement. Both these findings are likely a result of the short-lived increase in anabolism from *Pkm2* ablation. It has been suggested that PKM1 mediates ATP synthesis by promoting catabolism in the absence of PkM2.^23,24^ However, Rajala *et al*. recently demonstrated that the upregulation of PKM1 from *Pkm2* deletion is unable to match the catalytic activity of endogenous PkM2, suggesting that PKM1 cannot compensate for PkM2 deficiency.^24^ Bringing these findings together, it is possible that PKM1 upregulation interferes with the anabolic influx from PkM2 inhibition and promotes catabolism. The resultant inhibition of anabolism deprives photoreceptors of essential building blocks, leading to the observed decrease in ONL thickness. At the same time, PKM1 is unable to generate sufficient energy to sustain photoreceptor function, resulting in the decrease in ERG response over time. Alternatively, it is likely that retinal degeneration impacts a wide range of independent metabolic pathways. As such, PkM2 ablation is only able to affect PkM2-dependent processes, leading to a partial but not complete rescue of the deleterious phenotypes.

Although this study identifies PkM2 as one of the major players in slowing down retinal degeneration, future studies identifying specific molecules and pathways would be beneficial in understanding the mechanism of this relationship. Wubben *et al*. offered critical insight into the specific genes impacted by PkM2 ablation in the retina.^23^ Expanding their analysis to include gene expression changes in the RPE offers a quantitative way to not only support *Pkm2*’s role in regulating metabolic coupling, but also track the metabolic changes over time. Another promising direction for research is to characterize PKM1’s role and whether its interaction with PkM2 is conserved in inherited retinal disease. This question is important as it would clarify whether PKM1 indeed acts as an antagonistic regulator of anabolic influx that may limit the impact of metabolic reprogramming. Other potential directions include tuberous sclerosis complex 1 (TSC1), an mTOR inhibitor, and hypoxia inducible factor-1α (HIF-1α), a transcription factor responsible for oxygen consumption regulation. In our earlier studies, we showed that both of these molecules are linked to photoreceptor rescue from metabolic reprogramming.^19,46^ Moreover, separate groups have found that PkM2 interacts directly with mTOR and HIF-1α, hinting towards possible mechanistic pathways.^47–49^

One important consideration is that Rajala *et al*. have shown *in vitro* evidence that PkM2 may regulate transcriptional activity through the PDE6β promoter.^24^ This finding suggests that PKM2 knockout may have promoted photoreceptor health by working directly with Pde6β, which would indicate that the beneficial effects are specific to Pde6β deficiency. As such, we highlight the need to use a different RP model to evaluate whether the rescue phenotypes from PKM2 ablation are in fact a result of its role in a gene non-specific metabolic pathway.

Current CRISPR/Cas9-based gene therapy relies on a gene-and mutation-specific approach. Given that over 80 genes have been implicated in the development of RP, this strategy is impractical as it is not amenable to large-scale production or treatment. Here, we have demonstrated that PKM2 ablation offers 6 equivalent years of improved eyesight in our RP model,^50^ suggesting that elevations in rod-specific anabolism can offer protection against degeneration in a non-gene-specific manner. Importantly, the metabolic pathways highlighted in this study are common to not only retinal degeneration, but also neurodegenerative conditions including Alzheimer’s Disease.^51,52^ As such, reprogramming the metabolism of a terminally differentiated neuron towards a more anabolic state may offer a universal, gene non-specific treatment for multiple neurodegenerative conditions. The classical function of PKM2 as a catalyst for the final step in glycolysis lies at the heart of metabolic regulation. Therefore, a deeper understanding of its role in coupling the dynamic metabolisms of the retina and RPE may offer critical insight into and advancement towards the development of gene non-specific therapeutics. Such a strategy can then be used synergistically with gene supplementation therapy to optimize efficacy.

## MATERIALS AND METHODS Animals

All mice were maintained in the Columbia University Pathogen-free Eye Institute Annex Animal Care Services Facility under a 12-hour light/12-hour dark cycle. *PKM2_flox/flox_* mice (B6;129S-*Pkm_tmi.iMgvh_*/J) were purchased from The Jackson Laboratory while *Pde6b_H620Q/H620Q_* mice were re-derived via oviduct transfer from morulae provided by the European Mouse Mutant Archive (EMMA). Our lab generated *Pde6g_CreERT2_* mice.^53^

The inducible *Pde6g*_CreERT2_ driver was used to exert both temporal and spatial control over PKM2 ablation in *Pde6β_H620Q/H620Q_* mice. PKM2 ablation was activated during the mid-stage of degeneration for clinical relevance and limited to the rod photoreceptors to prevent embryonic lethality. To generate this conditional PKM2 knockout in a *Pde6β_H620Q/H620Q_* mouse model, *Pde6β_H620Q/+_* mice were crossed first with *Pde6g_CreERT2_* mice and then *PKM2*_flox/flox_ mice. The resultant progeny were then backcrossed four generations to generate heterozygous *Pde6β_H620Q/+_ PKM2*_flox/flox_ *Pde6g*_CreERT2_. A total of six generations of backcrosses were performed to produce the desired homozygous genotype of *Pde6β_H620Q/H620Q_ Pkm2*_flox/flox_ *Pde6g*_CreERT2_.

The experimental mice were injected intraperitoneally with tamoxifen (amount: 100 μg/g body weight; concentration: 100 mg/ml in ethanol; dilution: 10 mg/ml in corn oil at 42 °C; catalog T5648; Sigma-Aldrich) at P8-11 to generate a genotype of *Pde6β_H620Q/H620Q_ Pkm2*_-/-_. Isogenic control and wild-type mice were injected with ethanol (amount: 100 μg/g body weight; dilution: 10% in corn oil) to generate a genotype of *Pde6β_H620Q/H620Q_ Pkm2*_+/+_. The mice were not discriminated based on sex.

All experiments were approved by the Columbia University Institutional Animal Care and Use Committee (IACUC) prior to the start of the study. All mice were used and maintained in accordance with the Statement for the Use of Animals in Ophthalmic and Vision Research of the Association for Research in Vision and Ophthalmology and the Policy for the Use of Animals in Neuroscience Research of the Society for Neuroscience.

### Immunoblotting

Following the protocol described by Tsang, *et al*.,^54,55^ retina samples were collected from mice two weeks after injection. To evaluate PKM2 expression levels, the samples were homogenized using a solution consisting of M-PER Mammalian Protein Extraction Reagent (Prod #78501 Thermo Scientific), phosphatase inhibitor cocktail (catalog P2850-5 mL; Sigma), and protease inhibitor cocktail (catalog P8340-1 mL; Sigma). After transfer onto nitrocellulose membrane (BioRad), samples were blocked in 5% skim milk 902887 MP Biomedicals, LLC). The membrane was then immersed overnight at 4 °C in the following primary antibodies: rabbit Anti-PKM2 (1:1000; #3198S; Cell Signaling Technology), mouse anti-β-actin (1:1000; ab125248; Abcam), and rabbit anti-phospho-PKM2 (1:1000; #3827S; Cell Signaling Technology). The following day, the membrane was washed in 0.5% PBST (500 μl Tween-20 in 1,000 ml PBS), and subsequently incubated in mouse monoclonal anti-rabbit IgG-HRP antibody (1:2000; ab99697; Abcam) and rabbit anti-mouse IgG-HRP antibody (1:2,000; sc-358914; Santa Cruz Biotechnology Inc.). Chemiluminescence (EMD Millipore) was detected using Biomax film (Kodak), and PKM2, P-PKM2, and actin levels were then quantified.

### Histology

As described by Velez, *et al*., Eyes were enucleated from euthanized mice, by following along the limbus to create a puncture at the 12 o’clock position. 56 Each eye was then placed in a solution of 1:2x Karnovsky fixative for 24 hours. Following fixation, eyes underwent a saline wash and then dehydration to be embedded in paraffin. Afterwards, hematoxylin and eosin were used to stain paraffin sections, which were then examined by light microscopy.

### Rod OS Measurement

Following euthanasia, the eye was enucleated followed by cornea and lens dissection and vitreous removal. At every 100 microns from the optic nerve, Rod OS was measured using a Image J (NIH, Bethesda)e. Measurements were taken for segments 500 microns away from the optic nerve on both eyes. All procedures were done in accordance with previously established IACUC guidelines.

### ERG

we obtained scotopic, mixed maximum, and photopic electroretinogram (ERG) recordings for experimental and control mice four and eight weeks after injection. Both eyes of each mouse (20 eyes) were analyzed and averaged due to the correlation between mouse eyes. Following a 12 hour overnight dark adaptation, Mice were anesthetized using the protocol described by Sancho-Pelluz, *et* al.^33^ All procedures performed on the mice were completed under dim lighting. Mice were dilated using a solution composed of phenylephrine hydrochloride (2.5%, Paragon) and tropicamide (1%, Akorn) that was administered topically. Mouse body temperature was maintained during all experiments using a heating pad. Three different experiments were performed – scotopic recording (pulse intensity: 0.001 Cd/m2), mixed maximal recording (pulse intensity: 3 Cd/m2), and photopic recording (pulse intensity: 30 Cd/m2). An ERG was conducted using Espion E2 Electroretinography System (Diagnosys LLC)]. All ERG data was collected using Espion E2 Electroretinography System (Diagnosys LLC).

### Statistical Analysis

Using the ERG results, the maximal b-wave wave of each eye was taken and used to fit a linear model using generalized least squares. This method of regression was chosen because of the correlation between the eyes of each mouse and to account for unequal variance between treated and non-treated mice.

## Acknowledgements

Jonas Children’s Vision Care and Bernard & Shirlee Brown Glaucoma Laboratory are supported by the National Institutes of Health [P30EY019007, R01EY018213, R01EY026682, R24EY027285, U01EY030580, National Cancer Institute Core [5P30CA013696], Foundation Fighting Blindness [TA-NMT-0116-0692-COLU], the Research to Prevent Blindness (RPB) Physician-Scientist Award, and unrestricted funds from RPB, New York, NY, USA. S.H.T. is a member of the RD-CURE Consortium and is supported by Kobi and Nancy Karp, the Crowley Family Fund, the Rosenbaum Family Foundation, the Tistou and Charlotte Kerstan Foundation, the Schneeweiss Stem Cell Fund, New York State [C029572], and the Gebroe Family Foundation. VBM is supported by NIH grants [R01EY024665, R01EY025225, R01EY024698, R21AG050437, P30EY026877], and RPB, New York, NY.

## Data Availability

All data in this manuscript are available for share.

## REFERENCES

1. Friedman DS, O’Colmain BJ, Munoz B, Tomany SC, McCarty C, de Jong PT, Nemesure B, Mitchell P, Kempen J. Prevalence of age-related macular degeneration in the United States. Arch Ophthalmol. 2004;122(4):564–572.

2. Schmier JK, Jones ML, Halpern MT. The burden of age-related macular degeneration. Pharmacoeconomics. 2006;24(4):319–334.

3. Punzo C, Kornacker K, Cepko CL. Stimulation of the insulin/mTOR pathway delays cone death in a mouse model of retinitis pigmentosa. Nat Neurosci. 2009;12(1):44–52.

4. Ferrari S, Di Iorio E, Barbaro V, Ponzin D, Sorrentino FS, Parmeggiani F. Retinitis pigmentosa: genes and disease mechanisms. Curr Genomics. 2011;12(4):238–249.

5. Campochiaro PA, Mir TA. The mechanism of cone cell death in Retinitis Pigmentosa. Prog Retin Eye Res. 2018;62:24–37.

6. da Cruz L, Dorn JD, Humayun MS, Dagnelie G, Handa J, Barale PO, Sahel JA, Stanga PE, Hafezi F, Safran AB, Salzmann J, Santos A, Birch D, Spencer R, Cideciyan AV, de Juan E, Duncan JL, Eliott D, Fawzi A, Olmos de Koo LC, Ho AC, Brown G, Haller J, Regillo C, Del Priore LV, Arditi A, Greenberg RJ, Argus IISG. Five-Year Safety and Performance Results from the Argus II Retinal Prosthesis System Clinical Trial. Ophthalmology. 2016;123(10):2248–2254.

7. da Cruz L, Coley BF, Dorn J, Merlini F, Filley E, Christopher P, Chen FK, Wuyyuru V, Sahel J, Stanga P, Humayun M, Greenberg RJ, Dagnelie G, Argus IISG. The Argus II epiretinal prosthesis system allows letter and word reading and long-term function in patients with profound vision loss. Br J Ophthalmol. 2013;97(5):632–636.

8. Maguire AM, Simonelli F, Pierce EA, Pugh EN, Jr., Mingozzi F, Bennicelli J, Banfi S, Marshall KA, Testa F, Surace EM, Rossi S, Lyubarsky A, Arruda VR, Konkle B, Stone E, Sun J, Jacobs J, Dell’Osso L, Hertle R, Ma JX, Redmond TM, Zhu X, Hauck B, Zelenaia O, Shindler KS, Maguire MG, Wright JF, Volpe NJ, McDonnell JW, Auricchio A, High KA, Bennett J. Safety and efficacy of gene transfer for Leber’s congenital amaurosis. N Engl J Med. 2008;358(21):2240–2248.

9. Bainbridge JW, Smith AJ, Barker SS, Robbie S, Henderson R, Balaggan K, Viswanathan A, Holder GE, Stockman A, Tyler N, Petersen-Jones S, Bhattacharya SS, Thrasher AJ, Fitzke FW, Carter BJ, Rubin GS, Moore AT, Ali RR. Effect of gene therapy on visual function in Leber’s congenital amaurosis. N Engl J Med. 2008;358(21):2231–2239.

10. Russell S, Bennett J, Wellman JA, Chung DC, Yu ZF, Tillman A, Wittes J, Pappas J, Elci O, McCague S, Cross D, Marshall KA, Walshire J, Kehoe TL, Reichert H, Davis M, Raffini L, George LA, Hudson FP, Dingfield L, Zhu X, Haller JA, Sohn EH, Mahajan VB, Pfeifer W, Weckmann M, Johnson C, Gewaily D, Drack A, Stone E, Wachtel K, Simonelli F, Leroy BP, Wright JF, High KA, Maguire AM. Efficacy and safety of voretigene neparvovec (AAV2-hRPE65v2) in patients with RPE65-mediated inherited retinal dystrophy: a randomised, controlled, open-label, phase 3 trial. Lancet. 2017;390(10097):849–860.

11. Duncan JL, Pierce EA, Laster AM, Daiger SP, Birch DG, Ash JD, Iannaccone A, Flannery JG, Sahel JA, Zack DJ, Zarbin MA, and the Foundation Fighting Blindness Scientific Advisory B. Inherited Retinal Degenerations: Current Landscape and Knowledge Gaps. Transl Vis Sci Technol. 2018;7(4):6.

12. Takahashi VKL, Takiuti JT, Jauregui R, Tsang SH. Gene therapy in inherited retinal degenerative diseases, a review. Ophthalmic Genet. 2018;39(5):560–568.

13. Chinskey ND, Besirli CG, Zacks DN. Retinal cell death and current strategies in retinal neuroprotection. Curr Opin Ophthalmol. 2014;25(3):228–233.

14. Zadro-Lamoureux LA, Zacks DN, Baker AN, Zheng QD, Hauswirth WW, Tsilfidis C. XIAP effects on retinal detachment-induced photoreceptor apoptosis [corrected]. Invest Ophthalmol Vis Sci. 2009;50(3):1448–1453.

15. Punzo C, Xiong W, Cepko CL. Loss of daylight vision in retinal degeneration: are oxidative stress and metabolic dysregulation to blame? J Biol Chem. 2012;287(3):1642–1648.

16. Usui S, Komeima K, Lee SY, Jo YJ, Ueno S, Rogers BS, Wu Z, Shen J, Lu L, Oveson BC, Rabinovitch PS, Campochiaro PA. Increased expression of catalase and superoxide dismutase 2 reduces cone cell death in retinitis pigmentosa. Mol Ther. 2009;17(5):778–786.

17. Xiong W, MacColl Garfinkel AE, Li Y, Benowitz LI, Cepko CL. NRF2 promotes neuronal survival in neurodegeneration and acute nerve damage. J Clin Invest. 2015;125(4):1433–1445.

18. Iadevaia V, Huo Y, Zhang Z, Foster LJ, Proud CG. Roles of the mammalian target of rapamycin, mTOR, in controlling ribosome biogenesis and protein synthesis. Biochem Soc Trans. 2012;40(1):168–172.

19. Zhang L, Justus S, Xu Y, Pluchenik T, Hsu CW, Yang J, Duong JK, Lin CS, Jia Y, Bassuk AG, Mahajan VB, Tsang SH. Reprogramming towards anabolism impedes degeneration in a preclinical model of retinitis pigmentosa. Hum Mol Genet. 2016;25(19):4244–4255.

20. Martinez-Pastor B, Mostoslavsky R. Sirtuins, metabolism, and cancer. Front Pharmacol. 2012;3:22.

21. Dong G, Mao Q, Xia W, Xu Y, Wang J, Xu L, Jiang F. PKM2 and cancer: The function of PKM2 beyond glycolysis. Oncol Lett. 2016; 11(3): 1980–1986.

22. Rajala RV, Rajala A, Kooker C, Wang Y, Anderson RE. The Warburg Effect Mediator Pyruvate Kinase M2 Expression and Regulation in the Retina. Sci Rep. 2016;6:37727.

23. Wubben TJ, Pawar M, Smith A, Toolan K, Hager H, Besirli CG. Photoreceptor metabolic reprogramming provides survival advantage in acute stress while causing chronic degeneration. Sci Rep. 2017;7(1):17863.

24. Rajala A, Wang Y, Brush RS, Tsantilas K, Jankowski CSR, Lindsay KJ, Linton JD, Hurley JB, Anderson RE, Rajala RVS. Pyruvate kinase M2 regulates photoreceptor structure, function, and viability. Cell Death Dis. 2018;9(2):240.

25. McLaughlin ME, Ehrhart TL, Berson EL, Dryja TP. Mutation spectrum of the gene encoding the beta subunit of rod phosphodiesterase among patients with autosomal recessive retinitis pigmentosa. Proc Natl Acad Sci U S A. 1995;92(8):3249–3253.

26. Bird AC. Retinal photoreceptor dystrophies: the LI. Edward Jackson Memorial Lecture. American Journal of Opthalmology. 1995;119:543–562.

27. Hartong DT, Berson EL, Dryja TP. Retinitis pigmentosa. Lancet. 2006;368(9549):1795–1809.

28. Daiger SP, Bowne SJ, Sullivan LS. Perspective on genes and mutations causing retinitis pigmentosa. Arch Ophthalmol. 2007;125(2): 151–158.

29. Davis RJ, Tosi J, Janisch KM, Kasanuki JM, Wang NK, Kong J, Tsui I, Cilluffo M, Woodruff ML, Fain GL, Lin CS, Tsang SH. Functional rescue of degenerating photoreceptors in mice homozygous for a hypomorphic cGMP phosphodiesterase 6 b allele (Pde6bH620Q). Invest Ophthalmol Vis Sci. 2008;49(11):5067–5076.

30. Tsang SH, Tsui I, Chou CL, Zernant J, Haamer E, Iranmanesh R, Tosi J, Allikmets R. A novel mutation and phenotypes in phosphodiesterase 6 deficiency. Am J Ophthalmol. 2008;146(5):780–788.

31. Woodruff ML, Janisch KM, Peshenko IV, Dizhoor AM, Tsang SH, Fain GL. Modulation of phosphodiesterase6 turnoff during background illumination in mouse rod photoreceptors. J Neurosci. 2008;28(9):2064–2074.

32. Janisch KM, Kasanuki JM, Naumann MC, Davis RJ, Lin CS, Semple-Rowland S, Tsang SH. Light-dependent phosphorylation of the gamma subunit of cGMP-phophodiesterase (PDE6gamma) at residue threonine 22 in intact photoreceptor neurons. Biochem Biophys Res Commun. 2009;390(4):1149–1153..

33. Sancho-Pelluz J, Cui X, Lee W, Tsai YT, Wu WH, Justus S, Washington I, Hsu CW, Park KS, Koch S, Velez G, Bassuk AG, Mahajan VB, Lin CS, Tsang SH. Mechanisms of neurodegeneration in a preclinical autosomal dominant retinitis pigmentosa knock-in model with a Rho(D190N) mutation. Cell Mol Life Sci. 2019;76(18):3657–3665.

34. Tosi J, Davis RJ, Wang NK, Naumann M, Lin CS, Tsang SH. shRNA knockdown of guanylate cyclase 2e or cyclic nucleotide gated channel alpha 1 increases photoreceptor survival in a cGMP phosphodiesterase mouse model of retinitis pigmentosa. J Cell Mol Med. 2011; 15(8): 1778–1787.

35. Tosi J, Sancho-Pelluz J, Davis RJ, Hsu CW, Wolpert KV, Sengillo JD, Lin CS, Tsang SH. Lentivirus-mediated expression of cDNA and shRNA slows degeneration in retinitis pigmentosa. Exp Biol Med (Maywood). 2011.

36. Wert KJ, Davis RJ, Sancho-Pelluz J, Nishina PM, Tsang SH. Gene therapy provides long-term visual function in a pre-clinical model of retinitis pigmentosa. Human molecular genetics. 2013;22(3):558–567.

37. Yang J, Naumann MC, Tsai YT, Tosi J, Erol D, Lin CS, Davis RJ, Tsang SH. Vigabatrin-induced retinal toxicity is partially mediated by signaling in rod and cone photoreceptors. Plos ONE. 2012;7(8): e43889. doi:10.1371/journal.pone.0043889.

38. Sohocki MM, Daiger SP, Bowne SJ, Rodriquez JA, Northrup H, Heckenlively JR, Birch DG, Mintz-Hittner H, Ruiz RS, Lewis RA, Saperstein DA, Sullivan LS. Prevalence of mutations causing retinitis pigmentosa and other inherited retinopathies. Hum Mutat. 2001;17(1):42–51.

39. Chinchore Y, Begaj T, Wu D, Drokhlyansky E, Cepko CL. Glycolytic reliance promotes anabolism in photoreceptors. Elife. 2017;6.

40. Venkatesh A, Ma S, Le YZ, Hall MN, Ruegg MA, Punzo C. Activated mTORC1 promotes long-term cone survival in retinitis pigmentosa mice. J Clin Invest. 2015;125(4):1446–1458.

41. Kanow MA, Giarmarco MM, Jankowski CS, Tsantilas K, Engel AL, Du J, Linton JD, Farnsworth CC, Sloat SR, Rountree A, Sweet IR, Lindsay KJ, Parker ED, Brockerhoff SE, Sadilek M, Chao JR, Hurley JB. Biochemical adaptations of the retina and retinal pigment epithelium support a metabolic ecosystem in the vertebrate eye. Elife. 2017;6.

42. Hamel C. Retinitis pigmentosa. Orphanet J Rare Dis. 2006;1:40.

43. Molday RS, Moritz OL. Photoreceptors at a glance. J Cell Sci. 2015;128(22):4039–4045.

44. Barabas P, Cutler Peck C, Krizaj D. Do calcium channel blockers rescue dying photoreceptors in the Pde6b (rd1) mouse? Adv Exp Med Biol. 2010;664:491–499.

45. Du J, Yanagida A, Knight K, Engel AL, Vo AH, Jankowski C, Sadilek M, Tran VT, Manson MA, Ramakrishnan A, Hurley JB, Chao JR. Reductive carboxylation is a major metabolic pathway in the retinal pigment epithelium. Proc Natl Acad Sci U S A. 2016; 113(51): 14710–14715.

46. Zhang L, Du J, Justus S, Hsu CW, Bonet-Ponce L, Wu WH, Tsai YT, Wu WP, Jia Y, Duong JK, Mahajan VB, Lin CS, Wang S, Hurley JB, Tsang SH. Reprogramming metabolism by targeting sirtuin 6 attenuates retinal degeneration. J Clin Invest. 2016;126(12):4659–4673.

47. Sun Q, Chen X, Ma J, Peng H, Wang F, Zha X, Wang Y, Jing Y, Yang H, Chen R, Chang L, Zhang Y, Goto J, Onda H, Chen T, Wang MR, Lu Y, You H, Kwiatkowski D, Zhang H. Mammalian target of rapamycin up-regulation of pyruvate kinase isoenzyme type M2 is critical for aerobic glycolysis and tumor growth. Proc Natl Acad Sci U S A. 2011;108(10):4129–4134.

48. Zhang Z, Deng X, Liu Y, Liu Y, Sun L, Chen F. PKM2, function and expression and regulation. Cell Biosci. 2019;9:52.

49. Anastasiou D, Poulogiannis G, Asara JM, Boxer MB, Jiang JK, Shen M, Bellinger G, Sasaki AT, Locasale JW, Auld DS, Thomas CJ, Vander Heiden MG, Cantley LC. Inhibition of pyruvate kinase M2 by reactive oxygen species contributes to cellular antioxidant responses. Science. 2011;334(6060):1278–1283.

50. Dutta S, Sengupta P. Men and mice: Relating their ages. Life Sci. 2016;152:244–248.

51. Demetrius LA, Driver J. Alzheimer’s as a metabolic disease. Biogerontology. 2013;14(6):641–649.

52. Demetrius LA, Simon DK. An inverse-Warburg effect and the origin of Alzheimer’s disease. Biogerontology. 2012;13(6):583–594.

53. Koch SF, Tsai YT, Duong JK, Wu WH, Hsu CW, Wu WP, Bonet-Ponce L, Lin CS, Tsang SH. Halting progressive neurodegeneration in advanced retinitis pigmentosa. J Clin Invest. 2015;125(9):3704–3713.

54. Tsang SH, Burns ME, Calvert PD, Gouras P, Baylor DA, Goff SP, Arshavsky VY. Role for the target enzyme in deactivation of photoreceptor G protein in vivo. Science. 1998;282(5386): 117–121.

55. Tsang SH, Gouras P, Yamashita CK, Kjeldbye H, Fisher J, Farber DB, Goff SP. Retinal degeneration in mice lacking the gamma subunit of the rod cGMP phosphodiesterase. Science. 1996;272(5264): 1026–1029.

56. Velez G, Tsang SH, Tsai YT, Hsu CW, Gore A, Abdelhakim AH, Mahajan M, Silverman RH, Sparrow JR, Bassuk AG, Mahajan VB. Gene Therapy Restores Mfrp and Corrects Axial Eye Length. Sci Rep. 2017;7(1): 16151.

